# Increasing spectral DCM flexibility and speed by leveraging Julia’s ModelingToolkit and automated differentiation

**DOI:** 10.1101/2023.10.27.564407

**Authors:** David Hofmann, Anthony G. Chesebro, Chris Rackauckas, Lilianne R. Mujica-Parodi, Karl J. Friston, Alan Edelman, Helmut H. Strey

## Abstract

Using neuroimaging and electrophysiological data to infer neural parameter estimations from theoretical circuits requires solving the inverse problem. Here, we provide a new Julia language package designed to i) compose complex dynamical models in a simple and modular way with ModelingToolkit.jl, ii) implement parameter fitting based on spectral dynamic causal modeling (sDCM) using the Laplace approximation, analogous to MATLAB implementation in SPM12, and iii) leverage Julia’s unique strengths to increase accuracy and speed by employing Automatic Differentiation during the fitting procedure. To illustrate the utility of our flexible modular approach, we provide a method to improve correction for fMRI scanner field strengths (1.5T, 3T, 7T) when fitting models to real data.

---

Answering fundamental questions in neuroscience usually requires abstracting from neuronal populations to circuits (K. J. Friston et al., 2014; Hahn et al., 2020; Lee et al., 2022; Stephan et al., 2008). Neuronal circuits form causal models of interactions between different regions of the brain. Estimating model parameters, such as effective connectivity between brain regions, further requires data fitting (K. J. Friston, 2011; K. J. Friston et al., 2014; Stephan et al., 2008). One powerful approach is Dynamic Causal Modeling (DCM), introduced as part of the Statistical Parametric Mapping (SPM) toolbox (K. J. Friston et al., 2014; Stephan et al., 2008). DCM provides methods and an associated pipeline for modeling interactions between neuronal populations at the cortical level, fitting models from local field potential (LFP) recordings (K. J. Friston et al., 2015; R. J. Moran, Jung, et al., 2011; R. J. Moran, Mallet, et al., 2011), electroencephalography (EEG) data (David et al., 2006), and functional MRI (fMRI) sequences (Bernal-Casas et al., 2017; Cha et al., 2016; Rupprechter et al., 2020; Seghier & Friston, 2013; Sladky et al., 2015; Tik et al., 2018). Neuroscience applications of DCM have provided insights into both healthy (Bukhari et al., 2022; Hahn et al., 2020; Lee et al., 2022; Sladky et al., 2022) and impaired (Zarghami et al., 2023) neuronal functioning, in response to task-based (Hahn et al., 2020; Lee et al., 2022; Sladky et al., 2022) and resting-state (K. J. Friston et al., 2014; Zarghami et al., 2023) stimuli. Furthermore, the underlying mathematical approach of DCM has been extended to applications outside of neuroscience that benefit from causal inference (Parr et al., 2021).

Broadly speaking, DCM is a procedure used to infer model parameters from data and perform model comparison and selection (Novelli et al., 2024; Penny et al., 2004; Stephan et al., 2007; Zeidman et al., 2022). Models are defined that should describe the underlying neuronal dynamics, typically at the meso- or macro-scale, as well as the measurement process (BOLD, EEG, LFP). A variational Bayes approach is used to estimate the posterior distribution of the parameters by optimizing the evidence lower bound (R. J. Moran, Jung, et al., 2011; Novelli et al., 2024; Zeidman et al., 2022). Finally, the evidence lower bound or negative free energy is used for model comparison (K. Friston et al., 2007; Razi et al., 2017). By changing the prior distribution, a user can include and exclude parameters, such as connection weights between regions. Additionally, through model comparison, the user can judge which regions are most likely connected and whether they are excitatory or inhibitory. Over time, this approach has evolved and become more powerful, including the adoption of a computationally more efficient DCM algorithm in the spectral domain (K. J. Friston et al., 2014). Signals, assumed to be stationary, are transformed to the spectral domain by the computation of cross-spectral densities (CSD). This computation is performed by fitting an autoregressive (AR) model to the time series (in SPM12 the AR model is of fixed order 8). From the parameters of the AR model one can analytically calculate the CSD. In our package the fitting of the AR model is performed by standard maximum likelihood. Note that the standard procedure in SPM12 is a Bayesian fit defined in *spm mar.m*.

The most current implementation of DCM, as part of the Statistical Parametric Mapping toolbox (SPM12; available at fil.ion.ucl.ac.uk/spm/software/spm12/), has been widely and successfully applied to a variety of data modalities, cognitive functions, and disease processes (Hahn et al., 2020; Lee et al., 2022; Sladky et al., 2022; Zarghami et al., 2023). Here, we build on this achievement to extend its capabilities in two ways. First, one of the practical roadblocks to the wide adoption of multi-scale computational neuroscience has been limitations due to computation speed. DCM’s implementation in MATLAB introduces inherent computational barriers that become increasingly problematic as datasets scale to meet current reproducibility standards (Marek et al., 2022). For this reason, we chose to code our package in Julia, as part of the Neuroblox computational neuroscience platform (neuroblox.org), as Julia is a high-performance computing language that is known to show orders of magnitude speed improvements in ODE systems compared to both MATLAB and Python (Rackauckas, 2023). Second, we wanted to incorporate additional flexibility into the pipeline in terms of easy access to choices for neural mass models, model assembly, transfer functions within and between scales and modalities, as well as parameter fitting procedures.

In this work, we provide a new implementation of spectral DCM in the Julia programming language (julialang.org), building on the ModelingToolkit package (Ma, Gowda, et al., 2021). We demonstrate that this implementation can offer results identical to the original SPM12 implementation at a fraction of the computation speed in an open-source language. Additionally, we construct this implementation to allow for great user flexibility in model selection and specification, opening the door to new modeling techniques not achieved in prior implementations.

## Spectral Dynamic Causal Modeling

The distinctive feature of spectral DCM as compared to other forms of DCM is that it models the cross-spectral density of a signal. This implies the assumption of stationarity of the covariance matrix of the signals during measurement Novelli et al., 2024. The usual continuous-time state-space formulation with a measurement function is assumed

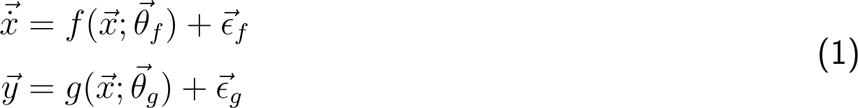

For Eqns. 1, 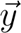 denotes the measurement of the (hidden) dynamics 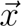, which could, for example, be the neuronal dynamics and hemodynamic response. 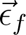 and 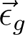 denote intrinsic or endogenous noise and measurement noise, respectively. In spectral DCM these are assumed to follow a power law distribution in frequency space. In fMRI measurements, the states *x* comprise not just the neuronal dynamics 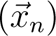 but also the hemodynamic response to the neuronal activity 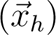. A further assumption made by spectral DCM is linearity of the hidden neuronal dynamics 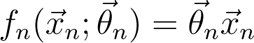 The dynamics for the hemodynamic response 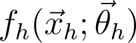 are modeled by the balloon model, and the measured signal is the blood oxygen level depletion (BOLD) denoted by 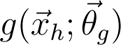. Both models are detailed in (Stephan et al., 2007), which is based on (Buxton et al., 2004). Here, we describe the model used in the code in more detail. The balloon model describes the dynamics of blood vessels in response to the neuronal signals. The cerebral blood flow *u*(*t*) is driven by the neuronal activity *x_n_* (for the sake of notational simplicity we omit an index that denotes a specific region of the multidimensional vector 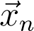 describing the neuronal activity of all regions. In what follows all dynamic variables relate to a single region.) through the induction of a vasodilatory signal *s*(*t*) governed by the following dynamics:

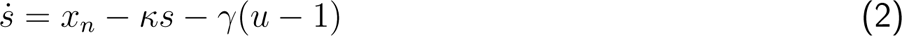

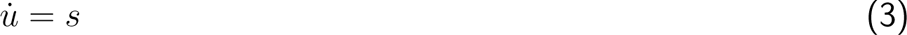

Moreover, the change in volume *v*(*t*) due to the blood flow as well as the related changes of the blood oxygenation level *q*(*t*) are given, according to (Buxton et al., 2004), by

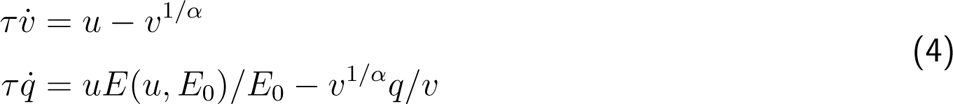

with *E* (*u,E* _0_) = 1−(1*−E* _0_)^1/u^ as a reasonable approximation across a wide range of conditions (Buxton & Frank, 1997; Stephan et al., 2007). Finally, based on blood oxygenation *q*(*t*) and blood volume *v*(*t*) we compute the BOLD signal according to the following equation

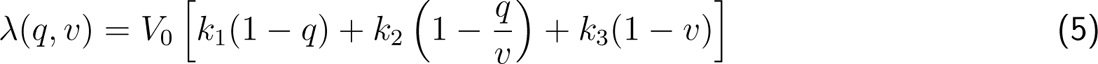

with the following parameter definitions

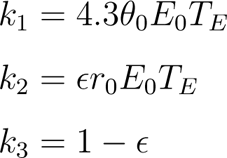

where ɛ is ratio of intra- and extravascular BOLD signal at rest, *E* _0_ is the resting oxygen extraction fraction, and *r*_0_ is the slope of the relation between the intravascular relaxation rate *R*\* and oxygen saturation. The solution to the differential equations Eqns. 1 can be approximated by the first order of a Taylor expansion as

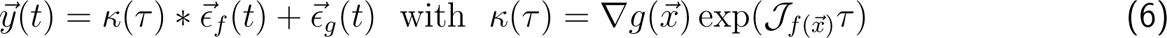

where 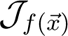 denotes the Jacobian of *f* w.r.t. 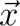 in Eqn. 1. To find an expression for the cross-spectral density we Fourier transform Eqn. 6

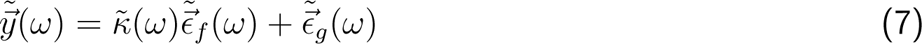

where 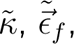, and 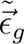 denote the Fourier transforms of the convolution kernel and noises respectively. The cross-spectral density is then

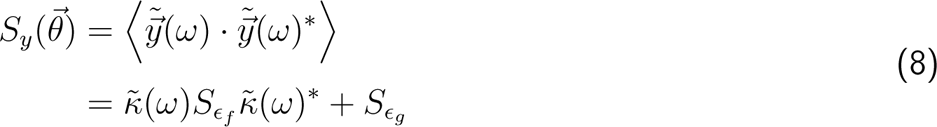

where * denotes the complex conjugate and 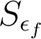 and 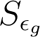 are the cross-spectral densities of the state noise and measurement noise, respectively. Note that, due to our assumption of uncorrelated noise, these are diagonal matrices. While the cross-spectrum of the measurement noise will have identical values on the diagonal, the state noise is composed of a global and a region-specific component, and thus diagonal elements can vary. This deviates from the presentation in (K. J. Friston et al., 2014) but is consistent with the implementation in SPM12. Note further that this derivation of the CSD implies that the underlying model is expanded at a fixed point solution in Eqn. 6, i.e. the model cannot be in a limit cycle or another dynamical state Bastos et al., 2015; R. Moran et al., 2008

We thus derived a linear approximation of the cross-spectral density 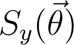 that is based on the details of the time-continuous state space model given in Eqns. 1. Next we would like to estimate the parameters of the model as well as the noise parameters and therefor need to define an optimization procedure. We follow the approach pursued by DCM, that is a Variational Bayes approach with Laplace approximation, termed Variational Laplace for brevity (Zeidman et al., 2022).

## Variational Laplace

An inverse problem consists of estimating parameters from data. In a Bayesian setting this implies computing the posterior distribution

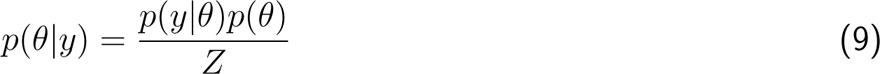

In the ideal setting, the true posterior distribution can analytically be computed from the likelihood *p*(*y|θ*), the prior distribution *p*(*θ*) and the normalization *Z*, also called the model evidence or marginalized likelihood 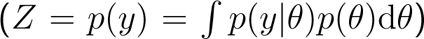 from the Bayes formula (Eqn. 9). However, typically, there is no such closed-form solution, and instead, we have to resort to sampling techniques (e.g., Markov Chain Monte Carlo), which are unbiased estimators, or we compute an approximation to the posterior. Variational Bayes is such an approximation approach that borrows from variational calculus by introducing a parameterized function *q*(*θ*) that will be shaped, through an optimization procedure, to approximate the true posterior *q*(*θ*) *≈ p*(*θ|y*). This optimization procedure requires the definition of an objective function which is typically the evidence lower bound (ELBO), sometimes also called the variational lower bound or free energy (particularly in the context of DCM) due to its resemblance of the functional form of the free energy in physics (Zeidman et al., 2022).

There are different ways to derive the ELBO; one approach is to start from the Kullback-Leibler (KL) divergence between the variational distribution and the posterior *D* _KL_(*q*(*θ*)*||p*(*θ|y*)) (Cover & Thomas, 2006). Minimizing *D* _KL_ is equivalent to maximizing the ELBO:

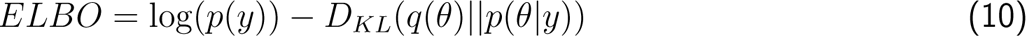

Note that the KL divergence is only well defined if *q* has probability 0 at the same intervals as *p*; in other words, the support of the two distributions must be the same.

Applying the Laplace approximation to simplify this problem means assuming a Gaussian variational posterior *q*(*θ*) = *N* (*µ*_θ_, Σ_θ_), hence the term variational Laplace. This is the approach taken by DCM (Novelli et al., 2024).

Here, we present several advances that expand the capabilities of spectral DCM as implemented in SPM12. We provide a toolbox with a user-friendly interface for adding new models and linking them in arbitrarily complex circuits. The toolbox also includes built-in corrections for different acquisition field strengths not previously considered in DCM analyses. This software leverages ModelingToolkit.jl, a symbolic computation framework written in the programming language Julia, to achieve a high degree of modularity and thereby easy model composability (Fig. 1). While the implementation of DCM in Julia already provides a speed increase over the traditional MATLAB version, we also employ automatic differentiation to further improve computation speed.

**Figure 1:**
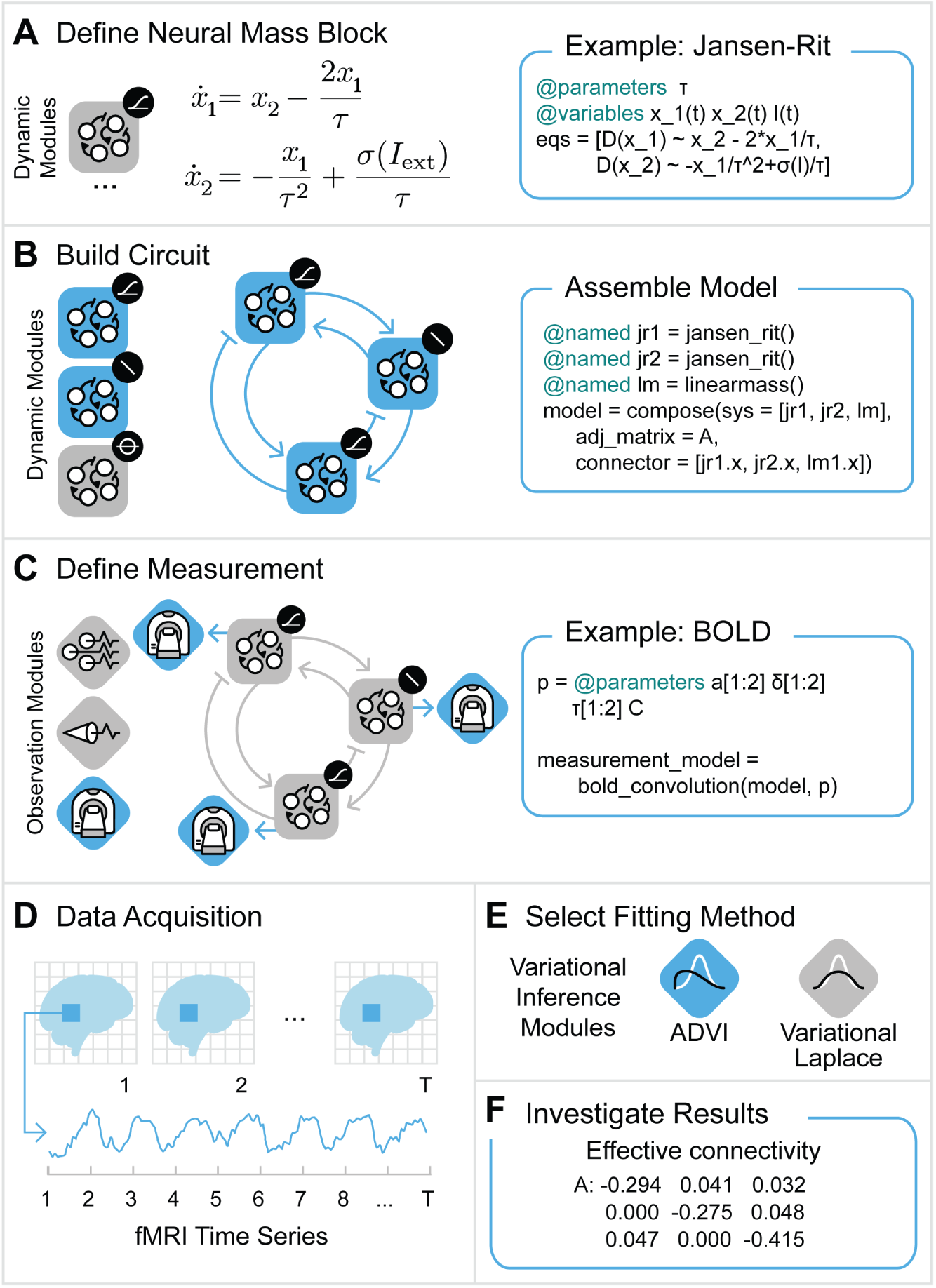
Modular design for user-friendly circuit model assembly and data fitting. Blue boxes on the right side in panels **A** to **C** show code snippets of the model assembly illustrated on the left. Note that the blue icon color just illustrates selection in the respective panel and carries no additional meaning. In **A** the definition of a model (also called block) is shown, in this case a Jansen-Rit model. Coding it is almost as simple as writing its differential equations. A user can pick and choose from a library of available models (panel **B** on the very left illustrates a few choices disambiguated by the small black icon on the top-right corner of the large icon. From top to bottom: Jensen-Rit model, linear neural mass model, and next generation neural mass model. These blocks don’t exhaust available models for neuronal dynamics in our software Neuroblox. A linear neural mass and two Jansen-Rit blocks are selected and connected to form a circuit. In addition to a model for the neuronal activity, we select (or define a new) measurement modality in **C**; in this particular case, we assume fMRI measurements and thus add a BOLD signal model. For LFP modeling lead-field equations are available in the package. Every region can in principle be described by a different neural mass model which substantially expands modeling power over the SPM12 toolbox, where regions are constrained to have dynamics of the same functional form. Lastly, a fitting procedure can be selected (**E**) and parameter values estimated (**F**) from acquired data (**D**). (Graphic in (**D**) adopted from CC-licensed graphic in Sidén, 2020) differential equations (see Fig. 1A).

## Modular design for user-friendly circuit model assembly and data fitting

Defining a model should be as simple as writing down its equations. For this purpose our toolbox design leverages ModelingToolkit.jl (Ma, Gowda, et al., 2021) and Symbolics.jl (Gowda et al., 2022) for symbolic computation which provide a simple and intuitive way to define a model that consists of differential equations (see Fig. 1A).

For instance, the user can define arbitrary models describing the dynamics of a particular brain region and assemble them into an overall model through an adjacency matrix (see Fig. 1B). In this paper we studied two different systems for the neuronal dynamics: a simple linear ODE system as is typically used in SPM12 for the spectral DCM analysis of fMRI signals (K. J. Friston et al., 2014), and a more complex canonical micro-circuit model which is used to model LFP signals (Bastos et al., 2012; R. J. Moran, Jung, et al., 2011; R. J. Moran et al., 2013) (see Fig. 2). Definition of an observation model that relates the (hidden) neuronal activity to the actual neurophysiological measurement completes the model assembly (see Fig. 1C). By providing measurements corresponding to the observation modality (Fig. 1D) the user can then pick a variational inference algorithm, besides Variational Laplace that is employed by SPM12 and studied in this manuscript, we also provide Automatic Differentiation Variational Inference (ADVI) as implemented in the library Turing.jl (see Ge et al., 2018; Kucukelbir et al., 2016) (Fig. 1E) to estimate the posterior probability of the model parameters, for instance the effective connectivity between regions (Fig. 1F).

**Figure 2:**
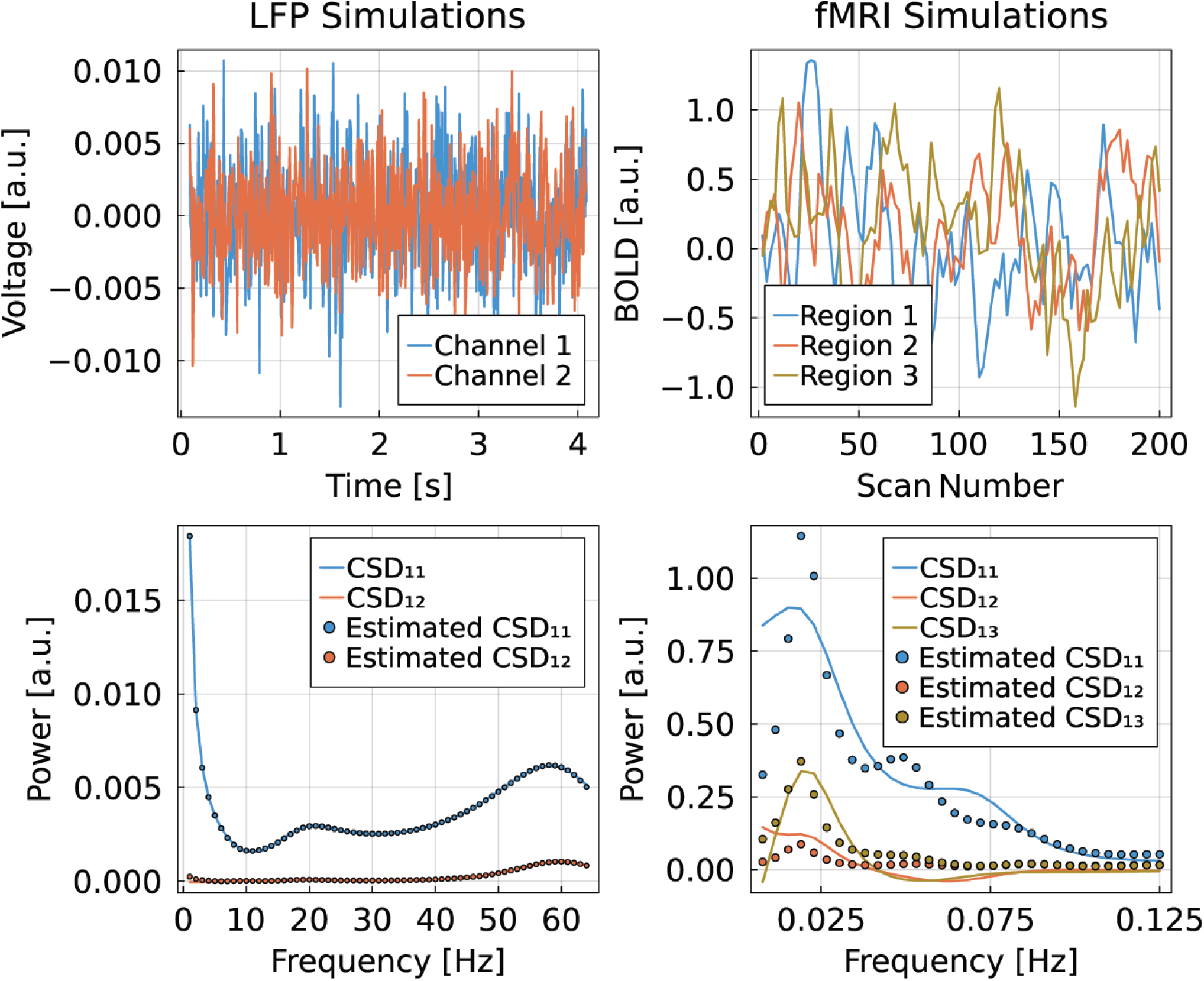
Estimation of cross-spectral densities of simulated data. Top row are examples of simulated signals, left are LFP measurements of two channels, and right are fMRI measurements of three regions. Plots in the bottom row are examples of simulated (lines) and estimated (dots) cross-spectral densities. We find a nearly perfect match for LFP signals, while fMRI signals show only a very rough agreement. The key difference is the nonlinear transformation of the neuronal signals through the hemodynamic response function; the LFP signals are only affected by a constant gain term that may vary for each channel. These simulations are based on the SPM12 scripts *DEM demo induced FMRI.m* for fMRI signals and *spm csd demo.m* for LFP signals. For details about the LFP model see R. J. Moran, Jung, et al., 2011

The toolbox provides a Variational Laplace inference algorithm analogous to the SPM12 implementation, as well as a Variational Laplace implementation that leverages automatic differentiation for the computation of the model Jacobian, both with and without symbolic computation. The inference is performed on the cross-spectral densities of the measured time series. The initial conditions of the states need to be selected so as to be in a steady state solution.

## Symbolic Computation

Using symbolic computation provides two important advantages. First, it facilitates modular definitions of models, which makes them easily extendable, alterable, and composable. Second, it provides a means of automatic analytical computation of gradients and Jacobians, which avoids the potentially imprecise numerical computation of derivatives or the effort of hard-coding the analytical derivatives for each new model. In spectral DCM, three multidimensional derivatives need to be computed. The gradient of the measurement function *g* and the Jacobian of *f* need to be calculated for the computation of the cross-spectral densities, both with respect to the system states (see Eqn. 6). Then, the Jacobian needs to be calculated with respect to the parameters for we wish to optimize, to compute the optimization update steps (K. J. Friston et al., 2014; Novelli et al., 2024; Zeidman et al., 2022). Symbolic computation is based on the packages ModlingToolkit.jl and Symbolics.jl (Gowda et al., 2022; Ma, Gowda, et al., 2021)

## Automatic Differentiation

Julia provides different libraries for performing automatic differentiation. We chose ForwardDiff.jl because of its robustness and good performance for systems with up to 100 dimensions (Ma, Dixit, et al., 2021; Revels et al., 2016).

The SPM12 implementation of the matrix-exponential of the kernel *κ* (*τ*) given in Eqn. 6 requires an eigenvalue decomposition. This poses a particular challenge since the eigenvalue decomposition in the standard library LinearAlgebra.jl is not a native Julia implementation but based on the external, compiled BLAS library and thereby is not amenable to automatic differentiation. The package GeneralizedLinear-Algebra.jl provides linear algebra operations that are fully implemented in Julia; however, its eigenvalue decomposition only provides eigenvalues but not the corresponding eigenvectors at the point of writing this manuscript.

Thus, we created a dispatch of the eigenvalue decomposition performed by the function *eigen* defined in LinearAlgebra.jl. Dynamic dispatch is the process of selecting which implementation of a polymorphic operation to call at run time. Polymorphic operation refers to the possibility to define different operations, in our case functions, that carry the same name but perform different operations. A dispatch of a function is a specific implementation of a function with an interface that differs in terms of number or types of arguments. This allows the programmer to define a particular behavior for particular types, like for instance the eigenvalue decomposition of a matrix of type Dual as is our case. The dispatch implements the analytical derivative of eigenvalues and eigenvectors (Lin et al., 2020; Van der Aa et al., 2007). However, the analytic derivative of eigenvectors with degenerate eigenvalues has poles and is thus numerically unstable around degeneracies. Indeed, the standard SPM initial conditions for the linear model and balloon model, which are by default set to zero, are resulting in a Jacobian with degenerate eigenvalues. To avoid this we introduce small Gaussian perturbations (of the order *O*(10*^−^*^2^)) to the initial conditions as well as to the prior expectation value of the connection matrix. This suffices to avoid this issue and does not affect the parameter estimations detrimentally. We note further that this procedure to avoid degeneracies depends on the specific model. For instance it is not required for coupled canonical microcircuits. Under such circumstances we keep the SPM12 standard and set all states to be 0, that is to the trivial fixed point of the model.

Complex numbers pose an additional hurdle since the Dual type of ForwardDiff.jl is not defined for complex numbers. This was simply overcome by dealing with the real and imaginary parts separately and then assembling the Dual of the real part and the Dual of the imaginary part into the complex number type. We improved in speed performance over a previous implementation that addresses these problems (Thummerer & Mikelsons, 2023).

## Julia implementation provides substantial speed improvements

Computational complexity can pose a bottleneck when analyzing high-dimensional datasets covering multiple brain regions. Julia provides a high-performance computing environment that reaches computational speeds comparable to C/C++ due to just-in-time compilation (Bezanson et al., 2018; Pelenitsyn et al., 2021). We implemented the variational inference method based on the Laplace approximation, also called Variational Laplace, for parameter estimation analogous to the implementation in SPM12. We compare our method based on simulated toy data with the SPM12 implementation to ensure consistency and correctness of our implementation in Julia.

We find that the Julia implementation provides a meaningful speed improvement (Fig. 3) while yielding the exact same results in terms of posterior means as well as free energy for all the different implemented versions.

**Figure 3:**
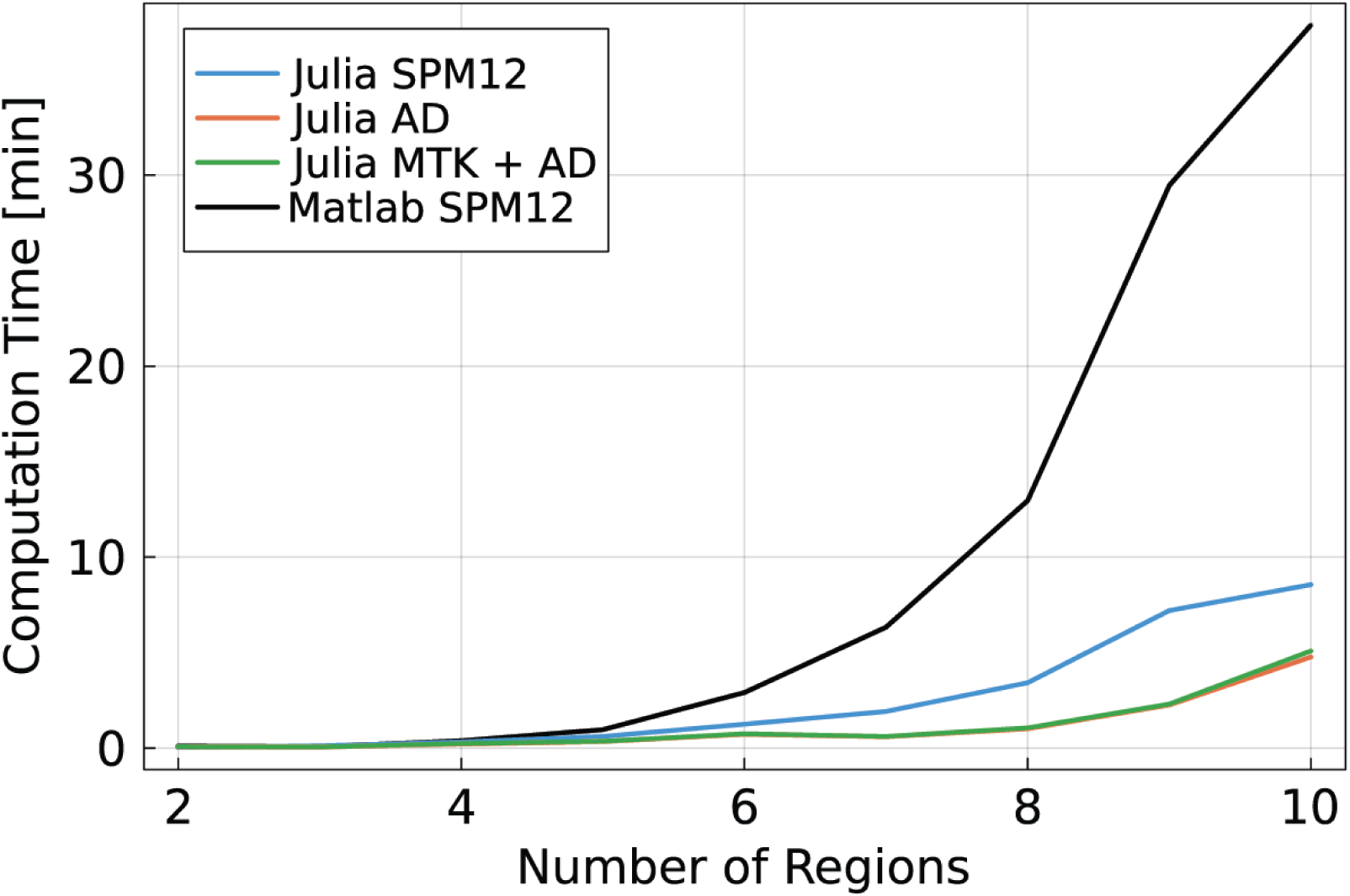
Julia implementation provides substantial speed improvements. Graphs show the speed increase with respect to the MATLAB SPM12 (black) implementation in dependence on the number of regions. Notable speed improvement is already achieved by a plain Julia re-implementation of the SPM12 DCM (blue). Employing Automatic Differentiation (AD) results in an additional speed improvement of up to 10-fold for 10 regions (orange). When additionally employing symbolic computation with ModelingToolkit.jl (MTK) to ensure compositionality of model building, computation speed suffers by a negligible factor as can be appreciated by observing the green line. This stems from an overhead introduced by symbolic computation that is specific to the current setup of the used libraries. However, the gain in flexibility by computing symbolic derivatives of arbitrary models over hard-coding them each time a new model is employed outweighs the subtle speed decrease. The alternative of numerically computing the gradient, which is done with SPM12, increases computational load and can suffer from inaccuracies.

These results were achieved using simulated data from the SPM12 script *DEM demo induced FMRI.m* . However, we note that when fitting coupled canonical microcircuits to simulated LFP data, we find that very small deviations (of order *O*(10*^−^*^3^) to *O*(10*^−^*^4^)) in parameter expectation value and free energy between the SPM12 implementation and the Julia version that makes use of AD and symbolic computation can occur (see Tab. 1). The simulation for this analysis is perfomed with the script *spm csd demo.m*.

**Table 1:** Comparison of estimated parameter expectations between SPM12 and Julia for simulations of the canonical microcircuit, based on the SPM12 script *spm csd demo.m*. We compare with the implementation that makes use of symbolic computation and automatic differentiation.

|  | SPM12 | Julia | ground truth |
| --- | --- | --- | --- |
| $a_{dp \rightarrow sp}$ | 0.4181 | 0.4181 | 1.0645 |
| $a_{dp \rightarrow ii}$ | 10.7195 | 10.7195 | 20.0855 |
| $a_{sp \rightarrow ss}$ | 0.9237 | 0.9237 | 1.6487 |
| $a_{sp \rightarrow dp}$ | 1.1089 | 1.1089 | 4.4817 |
| $L_1$ | 0.9797 | 0.9799 | 1 |
| $L_2$ | 0.8718 | 0.8719 | 1 |
| $r$ | 1.0567 | 1.0567 | 1 |
| $F$ | 1817.1634 | 1817.1660 | - |

Note that in SPM12 we need to set the DCM option “maxnodes” to a value equal to or larger than the number of regions we are inferring to ensure a correct comparison. Maxnodes will otherwise reduce the dimensionality of the problem to the value provided, which leads to a lower computational load.

Finally, we made the code amenable to automatic differentiation and use Julia’s ForwardDiff.jl library to perform automatic differentiation on the Variational Laplace optimization. We find a further speed improvement as can be observed in Fig. 3, yielding an almost order of magnitude speed improvement for 10 regions over the MATLAB SPM12 implementation.

In Dynamic Causal Modeling parameter fitting is the first step typically followed by model selection. This can be performed with our package in the same way as is performed in SPM12. By changing the prior expectation values for instance one can select effective connections to be present or absent and thereby changing model structure or complexity. Comparison of different models is performed by assessing their negative free energy values (see Tab. 2)

**Table 2:** By changing the prior mean to either 0 or 1 for the respective effective connection parameters, we perform a demonstration of DCM model selection. *F* _0_ is the free energy of the prior density and *F* is the free energy after the optimization procedure, as is typically output by DCM. A higher value indicates a better model fit to the date. As above, we perform the simulation of the canonical microcircuit with the SPM12 script *spm csd demo.m*. The true underlying connection is given in the bottom row and corresponds to the highest free energy value.

| $a_{dp \rightarrow sp}$ | $a_{dp \rightarrow ii}$ | $a_{sp \rightarrow ss}$ | $a_{sp \rightarrow dp}$ | $F - F_0$ |
| --- | --- | --- | --- | --- |
| 0 | 1 | 1 | 1 | 67 |
| 1 | 0 | 1 | 1 | 137 |
| 1 | 1 | 0 | 1 | 203 |
| 1 | 1 | 1 | 0 | 206 |
| 1 | 1 | 1 | 1 | 211 |

## fMRI pre-processing methods

As we are validating our pipeline against spectral DCM (K. J. Friston et al., 2014), we use a sample subject from the FC1000 project (available at www.fil.ion.ucl.ac.uk/spm/data/spDCM/) - one of the group used by Friston et al. (K. J. Friston et al., 2014) in their initial validation of spectral DCM - to demonstrate the coherence between SPM and Neuroblox implementations of DCM. Pre-processed data was available from this prior work, so this was the data used in the current study. Briefly, this scan was acquired at the University of Oxford Centre for Clinical Magnetic Resonance Research using a 3T Siemens scanner. Whole-brain fMRI was acquired using a gradient-echo EPI sequence (Repetition Time (*T* _R_) = 2000ms, Echo Time (*T* _E_) = 28ms, flip angle = 89,° field of view = 224mm, voxel size 3mm x 3mm x 3.5mm, 180 volumes acquired). This scan was accompanied by a T1-weigheted structural MRI (*T* _R_ = 2040ms, *T* _E_ = 4.7ms, flip angle = 8°, field of view = 192mm, voxel size = 1mm x 1mm x 1mm) to provide tissue segmentation. The fMRI scan was preprocessed by removing the first five volumes, realigning to the reference scan, normalizing to MNI space, correcting for slice-timing, and spatially smoothing the data with a Gaussian kernel (6mm FWHM). A GLM containing movement regressors was used to remove confounds, along with confound GLMs fit from the ventricle (CSF) and white matter signals (see SPM manual for details). Four ROIs were selected from the default mode network (DMN); all were chosen as 8mm radius spheres centered in each location. The four regions are the posterior cingulate cortex (PCC; center [0, -52, 26]), medial prefrontal cortex (mPFC; [3, 54, -2]), left intraparietal cortex (LIPC; [-50, -63, 32]) and right intraparietal cortex (RIPC; [48, -69, 35]).

## Comparison of SPM12 and Julia DCM fitting procedures on fMRI data

The DCM was fit in SPM12 using a standard pipeline. A fully connected DCM (i.e., all initial connection weights’ prior expectation value set to be non-zero, specifically they are set to one in SPM12) with no exogenous inputs was specified using the default parameter values (sequential data, *T* _E_ = 0.04, no modulatory effects, one state per region, no stochastic effects, no centering of input), then inverted using spectral DCM. Additionally, we probed the effects of field strength correction on the results by fitting at three different field strengths (1.5T, 3T, and 7T), presented in Tab. A2. Each field strength yielded significantly different results than the other, but all fits agreed in the directions of inter-regional connections, with the exception of LIPC intraregional connectivity (reversed at 1.5T).

### Discussion

DCM is one of the most widely used and validated methods to extract underlying neural connectivity from fMRI data by fitting and selecting models (Bernal-Casas et al., 2017; Cha et al., 2016; R. J. Moran, Mallet, et al., 2011; Novelli et al., 2024; Rupprechter et al., 2020; Seghier & Friston, 2013; Tik et al., 2018; Zarghami et al., 2023). The aim of the work presented here is two-fold: to improve the computational speed and accessibility of the well-established spectral DCM framework provided by SPM12 (K. J. Friston et al., 2014), and to increase the application domain by providing a modular framework for model fitting, thus empowering researchers to tailor their models to specific experimental setups and research questions. Importantly, each brain region can be defined to be governed by a different system of differential equations, which provides more flexibility compared to SPM12 where all regions are governed by the same system of differential equations (with potentially different parameters). We demonstrate that our implementation offers up to an order of magnitude improvement in computational speed compared to the SPM12 DCM, an enhancement enables efficient analysis of large-scale datasets and complex models.

Our work builds upon spectral DCM by re-implementing it in Julia, a high-performance programming language (Ma, Dixit, et al., 2021; Rackauckas, 2023). By implementing the variational Laplace approach presented here in Julia, we provide a method that recovers identical connectivity values - in both simulated and human fMRI data - as the SPM12 DCM approach while reducing computation time significantly. While SPM12 is implemented as open-source and easily accessible code, it relies upon Matlab, which inherently limits accessibility (to those who have a Matlab license) and computational speed (a known issue of Matlab code - see (Rackauckas, 2023)). Julia, on the other hand, has been built as open-source from the ground up, with a specific focus on optimizing for scientific computation (Bezanson et al., 2018; Danford et al., 2017; Freeman, 2015; Huang et al., 2019; Kramer et al., 2018; Ma, Dixit, et al., 2021). Even when fully parallelized Julia code offers order-of-magnitude speed benefits over similar Matlab implementations (Danford et al., 2017; Rackauckas, 2023). Although the translation of code into Julia is already sufficient to see a significant speed increase, we further optimize the approach by employing automatic differentiation (Revels et al., 2016). The adoption of automatic differentiation is also set to improve accuracy results for more complex non-linear models as presented in this study (Griewank & Walther, 2008).

A limitation of standard DCM implementation is that all but the most technically savvy users who feel comfortable editing the base code are constrained to the modeling choices imposed by the software. These prespecified models are a linear neural mass (attached to a balloon model for fMRI fitting (K. J. Friston et al., 2014)) or a few varieties of neuronal mass and neural field models for fitting LFP and EEG recordings (R. J. Moran, Jung, et al., 2011; R. J. Moran et al., 2013). In our implementation, we provide these standard approaches, but allow the user freedom to choose a variety of other neural mass models during fitting, or to specify their own choices. This is again made possible by implementation in Julia, specifically through the symbolic expressions offered in ModelingToolkit (Ma, Gowda, et al., 2021). As ModelingToolkit allows for symbolic expressions of neural masses, the process can be as simple as the Jansen-Rit example in Fig. 1A.

To illustrate the utility of a flexible modular approach, we provide a method to improve correction for MRI scanner field strengths when fitting a model to real data. Given the ubiquity and power of DCM approaches, it is necessary to consider advances in MRI scanner technology that have happened since the initial implementations of DCM, which used then state-of-the-art 1.5T and 3T scanners (K. J. Friston et al., 2014; Stephan et al., 2008). Here, we demonstrate how accounting for higher field strengths (e.g. 7T) that have become more common in human neuroimaging studies is an important consideration, as the results of fitting with the wrong field strength yield significantly different results. We note that the directionality of inter-regional connections is largely stable to this field strength correction (i.e., network topology does not change although the connection strengths do), giving us confidence that prior studies using this technique are still valid. However, given the noticeably different magnitudes of effective connectivity strengths, and the fact that this metric is used in many studies as the outcome of interest, it is important to apply the field strength of the original data when fitting a DCM, and we provide that capability in the package presented here.

The optimizations we provide here are only the beginning of future work in advanced neural modeling - indeed, this implementation is specifically built to encourage future development and open-sourced optimization. Some potential approaches to build on this include the substitution of alternatives to the variational Laplace optimization, such as ADVI (Kucukelbir et al., 2016) and NUTS sampler (Hoffman & Gelman, 2014) offered through Turing.jl (Ge et al., 2018). By being built within the Julia ecosystem, the method we present here is therefore poised for extensibility to accelerate the next generation of neural modeling.

## Author Contributions

Conceptualization: D.H., L.M.P. and H.H.S.; Formal analysis: D.H. and A.C.; Funding acquisition: C.R., A.E., L.M.P., and H.H.S.; Investigation: D.H.; Methodology: D.H.; Project administration: L.M.P.; Software: D.H., C.R., and H.H.S.; Supervision: C.R., K.J.F., and H.H.S.; Visualization: D.H. and A.C.; Writing – original draft: D.H. and A.C.; Writing - review & editing: D.H., A.C., L.M.P., K.J.F., and H.H.S.

## Acknowledgements

We thank Annabel Driussi for her valuable support with graphics. The research presented here was funded by the Baszucki Brain Research Fund, United States (LRMP). KJF is supported by funding for the Wellcome Centre for Human Neuroimaging (205103/Z/16/Z). AGC acknowledges the NIHGM MSTP Training Award, United States (T32-GM008444).

## Data and Code Availability

All methods shown here, which include an easy-to-use GUI as well as documentation and tutorials, are made freely available through the Neuroblox computational neuroscience platform (neuroblox.org). Neuroblox can be accessed through the JuliaHub repository (juliahub.com). The Julia code-base without GUI is available in this repository: https://github.com/Neuroblox/Spectral-Dynamic-Causal-Modeling.git

## Appendix

To further illustrate the flexibility of the implementation presented here, we highlight the importance of being able to fit fMRI data at different field strengths to match the field of acquisition. While this is possible in the traditional SPM12 implementation, it requires modifying the base code should field strengths other than the preset options be desired. As we illustrate in this Appendix, the field strength used to fit the data can significantly impact the results of the fit, and is therefore an important parameter for users to have easily accessible to best fit their data.

### Influence of field strength on hemodynamic variables in Balloon-Windkessel model

In the hemodynamic model DCM uses, six parameters can affect the fit substantially. Five of them have been discussed at length elsewhere (Buxton, 2013; K. J. Friston et al., 2003; Obata et al., 2004), and so will be briefly restated here. The echo time (*T* _E_) is always taken directly from the data being fit and is thus specified by individual users (K. J. Friston et al., 2014). The resting venous volume percent (*V*_0_) and resting oxygen extraction fraction (*E* _0_) are physiological parameters that - while variable across individuals - do not vary systematically in different field strengths and are thus assumed to be single values established in prior work (Buxton, 2013; K. J. Friston et al., 2003; Obata et al., 2004). The ratio of intra- to extravascular signal (ɛ) is fit on a region-by-region basis, which has been shown to improve accuracy (Stephan et al., 2007). The slope of intravascular relaxation rate as a function of oxygen (*r* _0_) can potentially vary at different field strengths; however, at fields above 3T, the contribution of this term becomes negligible due to the dominance of *k*_1_ in the balloon model (see Appendix A.3 in (Obata et al., 2004) for a full discussion of this effect). Given prior work that has demonstrated improved accuracy varying this parameter (Havlicek et al., 2015), however, we also vary it with *ν*_0_, as described below.

The remaining parameter of the balloon model is the maximum frequency offset at the outer surface of a magnetized vessel (*ν*_0_). This value has been shown to vary in different field strengths, and the exact derivation of this quantity has been addressed at length elsewhere. Summarizing this work (see (Buxton, 2013)): the maximum frequency offset at the surface of a blood vessel depends on the gyromagnetic ratio of protons in a field (*γ ≈* 128MHz), the susceptibility difference (Δ*χ* _0_ = 0.264 *×* 10*^−^*^6^), the hematocrit concentration (*H* = 0.44), the venous saturation (*Y* = 0.6), and the field strength *B* _0_, yielding the final expression of frequency offset

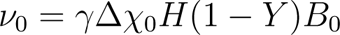

Prior work has estimated different values depending on different assumptions of hematocrit, venous saturation, or susceptibility difference, and individual variation (e.g., sex differences in baseline hematocrit concentration (Buxton, 2013)) can cause significant variability of this constant. However, for the work at hand, we note the linear dependence on field strength, with a doubling of this constant when moving from 1.5T to 3T, and even greater differences when moving to ultra-high fields (7T and 9.4T) used in modern human imaging studies. We probe these differences further in simulations and real data. In the following sections, the values for *ν*_0_ and *r* _0_ at different field strengths are taken from prior work (see (Havlicek et al., 2015) Tab. 1.B.). We do not vary the prior on ɛ purely for the sake of these illustrative examples, but there is no constraint on the user preventing them from doing so.

### Influence of field strength on hemodynamic response function

While there are many possible reasons for the effect that varying *ν*_0_ with field strength has on the fitting results of DCM, one important consideration is a direct effect on the shape of the hemodynamic response function (HRF). In this section, we probe the effect of field strength on the Balloon-Windkessel model HRF to determine what, if any, these effects are.

**Figure S1:**
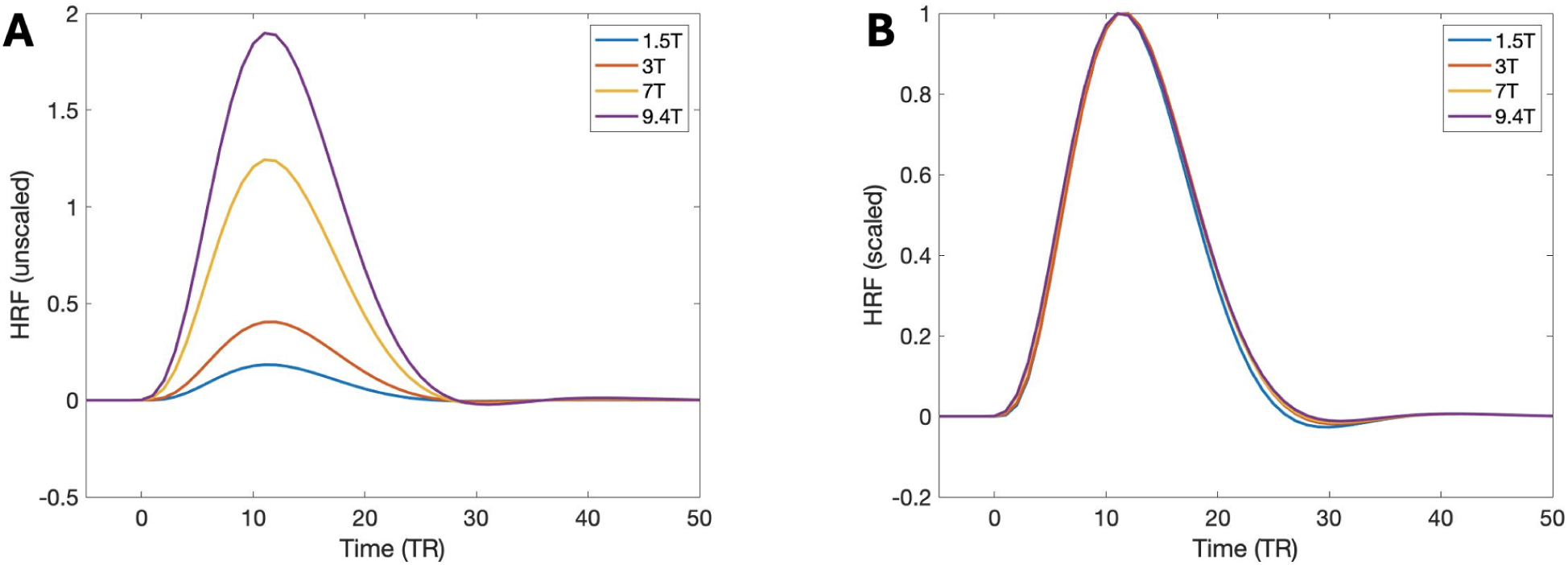
Field strength affects magnitude and shape of hemodynamic response function in Balloon-Windkessel model. A. The primary effect of field strength (captured by *ν*_0_) is the increase in magnitude of the HRF. The HRFs plotted here are the result of integrating the HRF in response to a Dirac delta pulse (simulated at TR=0.5s). B. To examine more subtle effects which remain after normalizing the fMRI signal, all field strengths from A are plotted on a normalized scale.

First, we must compute the HRF from the balloon model. As there are potential differences due to numerical solver effects between noisy simulations and pulsed simulations, we provide examples of both illustrating the effects of the varying field strength on the HRF. First, we compute the HRF in response to a Dirac delta pulse applied for a single TR. The results of these computations are shown in Fig. S1. Fig. S1A shows the strongest effect of field strength on the HRF - namely, the increase in magnitude of the response curve. However, as this can be removed by standard fMRI pre-processing techniques (signal normalization), Fig. S1B shows the same HRF curves normalized to a common scale. In this case, we observe a slight delay (*∼*250ms) in the peak times of higher fields (3T, 7T, and 9.4T) compared to the lowest field (1.5T). More importantly, the characteristic undershoot of the HRF takes on different depths depending on the field strength. Finally, it appears that the overall width of the peaks does not vary significantly.

To further probe this variability, we simulated the HRF model 50x at four different field strengths (1.5T, 3T, 7T, and 9.4T) common in human neuroimaging studies. Rather than using a simulated neuronal signal, we input white noise into the model to allow for computing the HRF directly from simulated data via division in the Fourier domain.

Considering the original neuronal signal *x* (*t*) and the output BOLD signal *y*(*t*), their relationship can be described by a convolution with an HRF *h*(*t*) such that

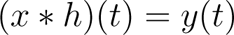

Converting this to the Fourier domain, this becomes a simple multiplication problem

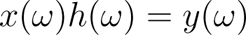

and the HRF can be determined by simple division if the input signal *x* (*t*) is white noise (guaranteeing *x* (*ω*) is a constant). We note that this approach does not work in most cases, as real signals can easily produce zero values in the Fourier domain, leading to a division by zero. However, in the case of pure noise as is considered here, we know the Fourier transform to be nearly constant with only minor variability, and so this approach is justified.

To further quantify the HRF differences across field strengths, we computed the mean peak and full width at half of the peak of each signal (results are shown in Tab. A1). We note that although there is a slight delay in the average time-to-peak of higher fields compared with 1.5T, this lies comfortably within the variance between simulations, and does not therefore make a strong contribution to variance in fits at different field strengths. Likewise, the duration of the HRF does not vary significantly across field strengths. This leads us to conclude that if the shape of the HRF does indeed play a role in goodness of DCM fits, it is in the more accurate representation of HRF undershoot at higher field strengths. We note, however, that there are still many questions regarding optimal fitting of the HRF at different field strengths that are left to a more complete discussion in future work.

**Table A1:** HRF characteristics at varied field strengths.

|  | 1.5T | 3T | 7T | 9.4T |
| --- | --- | --- | --- | --- |
| Time to peak (ms) | 6023 (506) | 6235 (390) | 6180 (476) | 6223 (575) |
| Width at half peak (ms) | 6323 (1091) | 6480 (1091) | 6283 (591) | 6289 (698) |

### Influence of field strength on hemodynamic response function

As field strength can impact the shape of the HRF, and can potentially impact model fitting as a consequence, we fit the same fMRI data presented in the previous section at three different field strengths: 1.5T, 3T, and 7T. As in the prior section, field strength was varied by varying *ν*_0_ and *r* _0_ according to prior work (Havlicek et al., 2015). The results of these fits are presented in Tab. A2 and visualized in Fig. S2.

**Table A2:** Fitting DCM from 3T rs-fMRI yields different results across field strengths. Results are presented as DCM fit (standard deviation).

| **1.5T** | PCC | mPFC | LIPC | RIPC |
| --- | --- | --- | --- | --- |
| PCC | 0.42 (0.088) | 0.19 (0.085) | 0.24 (0.059) | -0.03 (0.083) |
| mPFC | 0.05 (0.077) | 0.13 (0.094) | 0.06 (0.053) | 0.35 (0.070) |
| LIPC | 0.16 (0.080) | 0.56 (0.076) | 0.08 (0.087) | 0.05 (0.075) |
| RIPC | 0.19 (0.085) | -0.21 (0.084) | 0.23 (0.059) | 0.24 (0.081) |
| **3T** | PCC | mPFC | LIPC | RIPC |
| PCC | 0.58 (0.083) | 0.11 (0.092) | 0.34 (0.062) | -0.01 (0.090) |
| mPFC | 0.10 (0.087) | 0.50 (0.084) | 0.24 (0.060) | 0.18 (0.084) |
| LIPC | 0.11 (0.069) | 0.36 (0.069) | -0.01 (0.086) | 0.16 (0.070) |
| RIPC | 0.17 (0.089) | -0.03 (0.091) | 0.20 (0.063) | 0.50 (0.078) |
| **7T** | PCC | mPFC | LIPC | RIPC |
| PCC | 0.84 (0.075) | 0.14 (0.093) | 0.39 (0.065) | 0.09 (0.094) |
| mPFC | 0.15 (0.091) | 0.81 (0.076) | 0.35 (0.065) | 0.11 (0.092) |
| LIPC | 0.17 (0.075) | 0.30 (0.076) | 0.20 (0.082) | 0.15 (0.080) |
| RIPC | 0.12 (0.093) | 0.09 (0.093) | 0.26 (0.066) | 0.82 (0.073) |

To assess statistical significance of the differences across field strengths, we observe that the outputs of the DCM fits are Gaussian posterior distributions, and the difference between the fits can be computed as the difference of Gaussians where the mean is given as *µ*_diff_ = *|µ*_1_ *− µ*_2_*|* and the standard deviation as 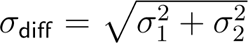 . Using this definition, we assess statistically significant differences as those whose difference distribution has a cumulative density function ¡0.005 at 0 (this is equivalent a one-tailed t-test p ¡ 0.005, or two-tailed t-test p ¡ 0.01). Turning to the results, we first note that the fits are significantly different across field strengths, with the variance of fit connection weights more than an order of magnitude smaller than the parameters themselves. Second, we note that these associations are not simple linear multiples based on the field strength but significantly impact the relative weights of the associations. For example, mPFC *→* LIPC estimated connection strength increases fourfold when fit at 3T compared to 1.5T (0.06 *→* 0.24), but increases more modestly when fit at 7T compared to 3T (0.24 *→* 0.35). Finally, we note that although these results are significantly different, when examining modern field strengths (3T and 7T), most directions of associations are the same (i.e., the signs of all connection weights agree), although more sign agreement is present between 1.5T and 3T than between 7T and either lower field strength. Thus, while fitting 7T data using 3T parameters is suboptimal, this approach will likely yield directionally accurate results, even if the relative weights are suboptimal. Taking these results together, we believe there is a compelling case that field strength should be accounted for when fitting fMRI data to provide the most accurate results.

**Figure S2:**
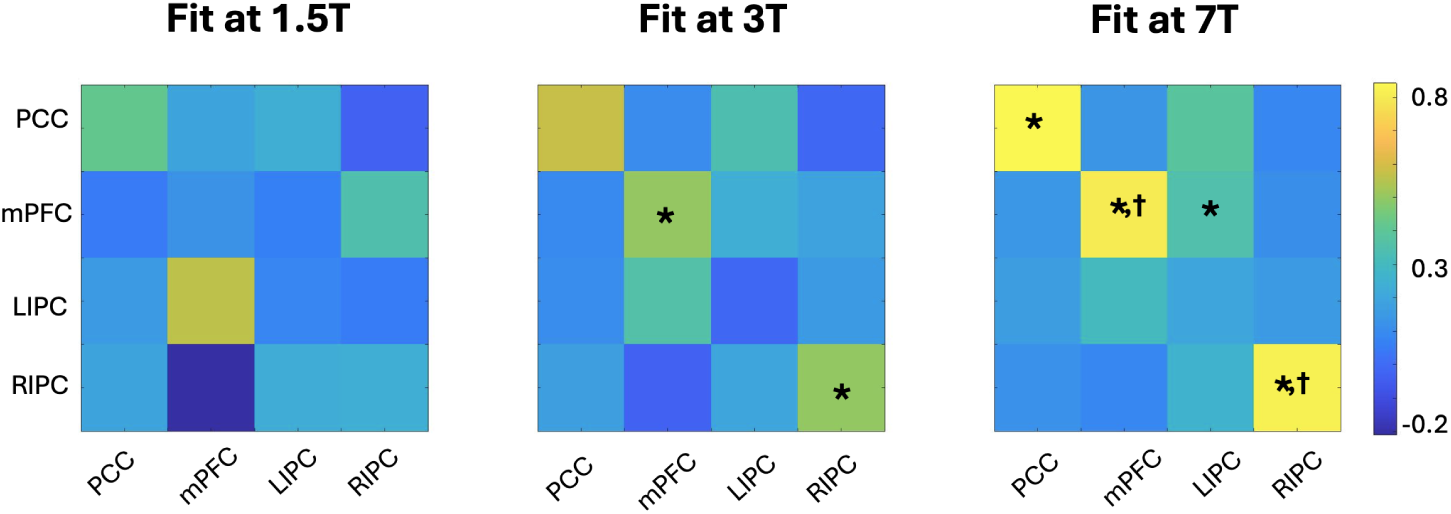
Visualization of connectivity strengths fit at different HRF field strength corrections. *Indicates significantly different than 1.5T fit. *†*Indicates significantly different than 3T fit. It is important to note that while the direction of all fits is the same in most cases, there are more differences in connectivity strengths, particularly noticeable at 7T, the current state of the art for clinical neuroimaging. These differences are especially pronounced compared to the fits done at 1.5T.

